# Admistration of Exogenous Melatonin Improves the Diurnal Rhythms of Gut Microbiota in High Fat Diet-Fed Mice

**DOI:** 10.1101/760421

**Authors:** Jie Yin, Yuying Li, Hui Han, Gang Liu, Xin Wu, Xingguo Huang, Rejun Fang, Kenkichi Baba, Peng Bin, Guoqiang Zhu, Wenkai Ren, Bie Tan, Gianluca Tosini, Xi He, Tiejun Li, Yulong Yin

## Abstract

Melatonin, a circadian hormone, has been reported to improve host lipid metabolism by reprogramming gut microbiota, which also exhibits rhythmicity in a light/dark cycle. However, the effect of admistartion of exogenous melatonin on the diurnal variation in gut microbiota in high fat diet (HFD)-fed mice is obscure. Here, we further confirmed the anti-obesogenic effect of melatonin on in mice feed with HFD for two weeks. Samples were collected every 4 h within a 24-h period and diurnal rhythms of clock genes expression (*Clock*, *Cry1*, *Cry2*, *Per1*, and *Per2*) and serum lipid indexes varied with diurnal time. Notably, *Clock* and triglycerides (TG) showed a marked rhythm only in the control and melatonin treated mice, but not in the HFD-fed mice. Rhythmicity of these parameters were similar between control and melatonin treated HFD mice compared with the HFD group, indicating an improvement of melatonin in the diurnal clock of host metabolism in HFD-fed mice. 16S rDNA sequencing showed that most microbiota exhibited a daily rhythmicity and the trends differentiated at different groups and different time points. We also identified several specific microbiota correlating with the circadian clock genes and serum lipid indexes, which might contribute the potential mechanism of melatonin in HFD-fed mice. Interestingly, administration of exogenous melatonin only at daytime exhibited higher resistance to HFD-induced lipid dysmetabolism than nighttime treatment companying with altered gut microbiota (*Lactobacillus*, *Intestinimonas*, and *Oscillibacter*). Importantly, the responses of microbiota transplanted mice to HFD feeding also varied at different transplanting times (8:00 and 16:00) and different microbiota donors. In summary, daily oscillations in the expression of circadian clock genes, serum lipid indexes, and gut microbiota, appears to be driven by a short-time feeding of an HFD. Administration of exogenous melatonin improved the compositions and diurnal rhythmicity of gut microbiota, which might be linked to host diurnal rhythm and metabolism.

**Importance:** Previous studies show that a circadian hormone, melatonin, involves in host lipid metabolism by reprogramming gut microbiota, which also exhibits rhythmicity in a light/dark cycle. However, the effect of melatonin drinking on the diurnal variation in gut microbiota in high fat diet-fed mice is obscure. Here, we found that 24-h oscillations were widely occurred in circadian clock genes, serum lipid indexes, and gut microbiota. Melatonin drinking improved the compositions and circadian rhythmicity of gut microbiota, which might be linked to host circadian rhythm and metabolism.

## 1. Introduction

Melatonin is a natural hormone mainly secreted by the pineal gland where its synthesis is driven by the master circadian clock located in the suprachiasmatic nucleous of the hypothalamus [1, 2]. Melatonin synthesis is activated by darkness and inhibited by light and thus this hormone is a key regulator of the circadian network [3–7]. In addition, melatonin also exerts various physiological functions (i.e., antioxidant function, bone formation, reproduction, cardiovascular, and immune regulation) and has been confirmed the therapeutic effects on gastrointestinal diseases, psychiatric disorders, cardiovascular diseases, and cancers [8–10]. More recently, a few studies have reported that melatonin receptor 1 knock-out mice show insulin and leptin resistance [11, 12], indicating a role of melatonin and its downstream signals in energy metabolism. Also, melatonin injection in lipopolysaccharide induced endotoxemia markedly improves energy metabolism by enhancing ATP production [13]. Similar potential of melatonin is also noticed in diabetes that lower melatonin secretion is independently associated with a higher risk of developing type 2 diabetes [14, 15]. These findings indicate an interation between melatonin signal and metabolic diseases. Indeed, Xu *et al.* also identified the anti-obese effect of melatonin in high fat diet (HFD)-induced obesity in a murine model by improving liver steatosis, low-grade inflammation, insulin resistance, and gut microbiota diversity and compositions [16]. We further confirmed the underlying mechanism of melatonin in HFD-induced lipid dysmetabolism which may be associated with reprogramming gut microbiota, especially, *Bacteroides* and *Alistipes*-mediated acetic acid production [17].

Gut microbiota is highly shaped by dietary HFD and obese human and animals are characterized by lower diversity and impaired gut microbiota compositions, especially for *Firmicutes* and *Bacteroidetes* abundances [18–21]. Interestingly, several reports have revealed that the gut microbiota and its metabolites exhibit circadian rhythm, which are driven by HFD [22–25]. Also, some microbe has been reported to be sensitive to melatonin [26], but the role of melatonin in the regulation of the diurnal patterns of gut microbial structure and function, and whether gut microbiota oscillations are associated with the anti-obese effect of melatonin is not yet known.

In this study we further analyzed the short-time effect of HFD feeding on diurnal variations in gut microbiota and the relationship between gut microbiota oscillations and the expression of circadian clock genes and serum lipids.

## 2. Materials and methods

### 2.1 Animal and diet

ICR mice were purchased from SLAC Laboratory Animal Central (Changsha, China). All animals had free access to food and drinking water (temperature, 25±2°C; relative humidity, 45-60%; lighting cycle, 12 h/d) during the experiment. Diets used in this study were according to our previous study [17]. As sex produces difference in melatoin profile [27], only female mice were used in this study to rule out the gender interference.

### 2.2 Melatonin treatment

126 Female mice (22.77 ± 0.10 g, about 4 week) were randomly grouped to control (Cont), HFD, and HFD plus melatonin (MelHF) groups (n=42). Mice in MelHF group received the HFD and melatonin water (0.4 mg/mL melatonin, Meilun, Dalian, China, directly diluted in drinking water) [17]. The melatonin solution was prepared daily and kept in a normal bottle with an aluminum foil cover to prevent light-induced degradation of melatonin. After two weeks of melatonin administration, 6 mice in each group were randomly killed at 0:00 (Zeitgeber time, ZT16), 4:00 (ZT20), 8:00 (ZT0, lights on), 12:00 (ZT4), 16:00 (ZT8), 20:00 (ZT12, lights off), and 24:00 (ZT16) O’clock (n=6). Blood samples were collected by orbital blooding. Liver, adipose tissues, and colonic digesta samples were weighed and collected.

### 2.3 Melatonin treatment at daytime and nighttime

Mice (26.89 ± 0.15 g) were randomly grouped to a control and three HFD groups (n=12). One group of HFD mice were received melatonin at daytime (8:00-16:00) and control water at night (16:00-8:00) (MelD) and another received melatonin at nighttime (16:00-8:00) and control water at daytime (8:00-16:00). All mice were killed at 8:00 am after 2-week feeding and samples were collected for further analyses.

### 2.4 Fecal microbiota transplantation

Mice were treated with antibiotics (1g/L streptomycin, 0.5g/L ampicillin, 1g/L gentamicin, and 0.5g/L vancomycin) to clear gut microbiota [17]. After 1-week of antibiotics treatment, the antibiotics-containing water was replaced with the regular water and the microbiota-depleted mice were transplanted with donor microbiota. Fecal supernatants from control (MT-Cont), HFD (MT-HF), and MelHF (MT-MelHF) (treated for 14 days) were transplanted into the microbiota-depleted mice at 8:00 and 16:00 (for 5 days). Following transplantation, all mice further received HFD and regular water for another 14 days.

### 2.5 Serum lipid indexes

Serum samples were separated after centrifugation at 3000 rpm for 10 min under 4 °C. Cobas c-311 coulter chemistry analyzer was used to test serum biochemical parameters, including triglycerides (TG), cholesterol (CHOL), high density lipoprotein (HDL), low density lipoprotein (LDL), glucose, and bile acid [17, 28].

### 2.6 RT-PCR

Total RNA from liver samples was isolated from liquid nitrogen frozen and ground tissues with TRIZOL regent (Invitrogen, USA) and then treated with DNase I (Invitrogen, USA). The reverse transcription was conducted at 37°C for 15 min, 95°C 5 sec. Primers used in this study were designed according to mouse sequence (Additional file 1: Supplementary table1). *β-actin* was chosen as the house-keeping gene to normalize target gene levels. The PCR cycling condition and the relative expression were according to previous studies [29–36].

### 2.7 Microbiota Profiling

Total genome DNA from colonic samples was extracted for amplification using specific primer with the barcode (16S V3+V4). Sequencing libraries were generated and analyzed according to our previous study [29, 37, 38]. OTUs were further used for genome prediction of microbial communities by Tax4Fun analysis [39].

### 2.8 Statistical Analysis

All statistical analyses were performed by using the one-way analysis of variance and multiple comparison was further conducted by Bonferroni analysis (SPSS 21 software). Correlation analysis was conducted by Pearson correlation analysis. The rhythmicity of clock genes, serum lipid indexes, and gut microbiota was assessed by cosinor analysis using the nonlinear regression model within Sigmaplot V 10.0 (Systat Software, San Jose, CA, USA) [40]. Data were expressed as the mean ± sem. A P value of < 0.05 was considered significant. All figures in this study were drawn by GraphPad Prism 7.04.

## 3. Results

### 3.1 Melatonin alleviates adipose accumulation in HFD-fed mice

Body weight was recorded in the current study and the results showed an increase in final body weight after 2-week HFD feeding (P<0.001) (Figure 1 A and B). Our previous study confirmed that administration of exogenous melatonin improved subcutaneous adipose accumulation in HFD-fed mice [17], the relative weight of subcutaneous adipose (P>0.05) tended be lower in the MelHF group. While visceral adipose tissue (P<0.05) were markedly reduced in the MelHF group in this study (Figure 1 C and D).

**Figure 1.**
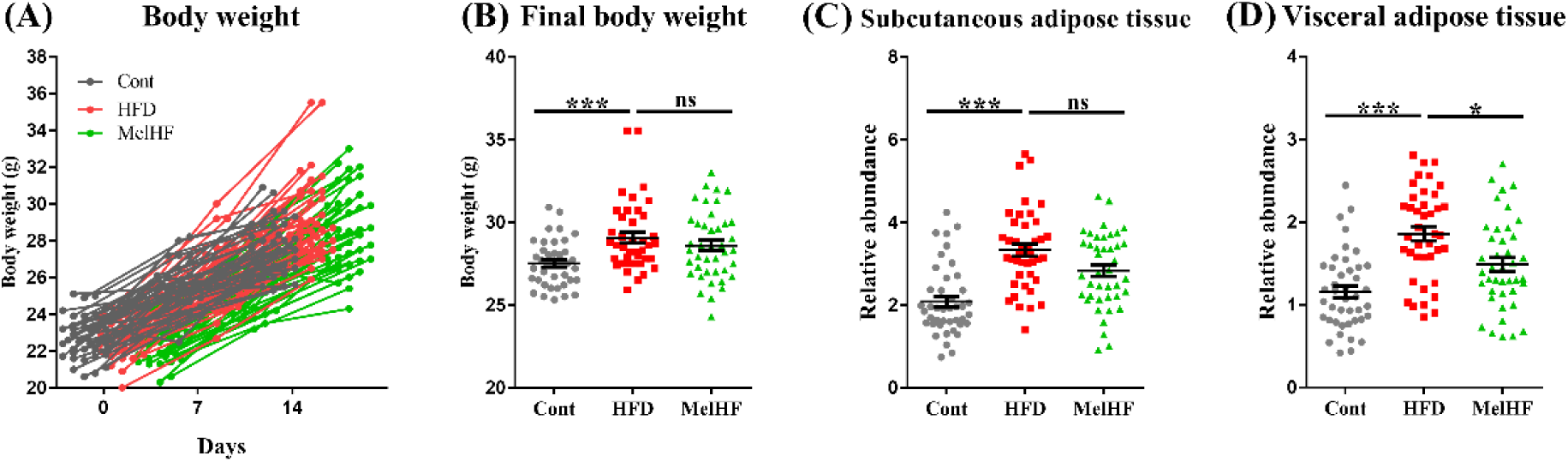
Effect of Melatonin treatment on body weight and lipid accumulation in HFD-fed mice. (A) Body weight (g), (B) final body weight (g), (C) relative weight of subcutaneous adipose tissue to body weight, and (D) relative weight of visceral adipose tissue to body weight (n=42). Values are presented as mean ± sem. Differences were assessed by Bonferroni test test and denoted as follows: *P<0.05; **P<0.01; ***P<0.001; ^ns^P>0.05.

### 3.2 Melatonin affectes clock gene expression in HFD-fed mice

Circadian clock and metabolism are generally impaired in HFD-fed mice [23, 41]. Thus, we further analyzed the diurnal variation of circadian clock genes (*Clock*, *Cry1*, *Cry2*, *Per1*, and *Per2*) in response to HFD and administration of exogenous melatonin (Figure 2 A; Table 1). Interestingly, *Clock* mRNA showed significant rhythmicity in the liver of control (P<0.01) and MelHF (P<0.05) mice, but not in HFD group (P>0.05), whereas the expressions of *Cry1*, *Cry2*, *Per1*, and *Per2* in the liver of showed a significant daily rhythm in all groups (P<0.05).

**Figure 2.**
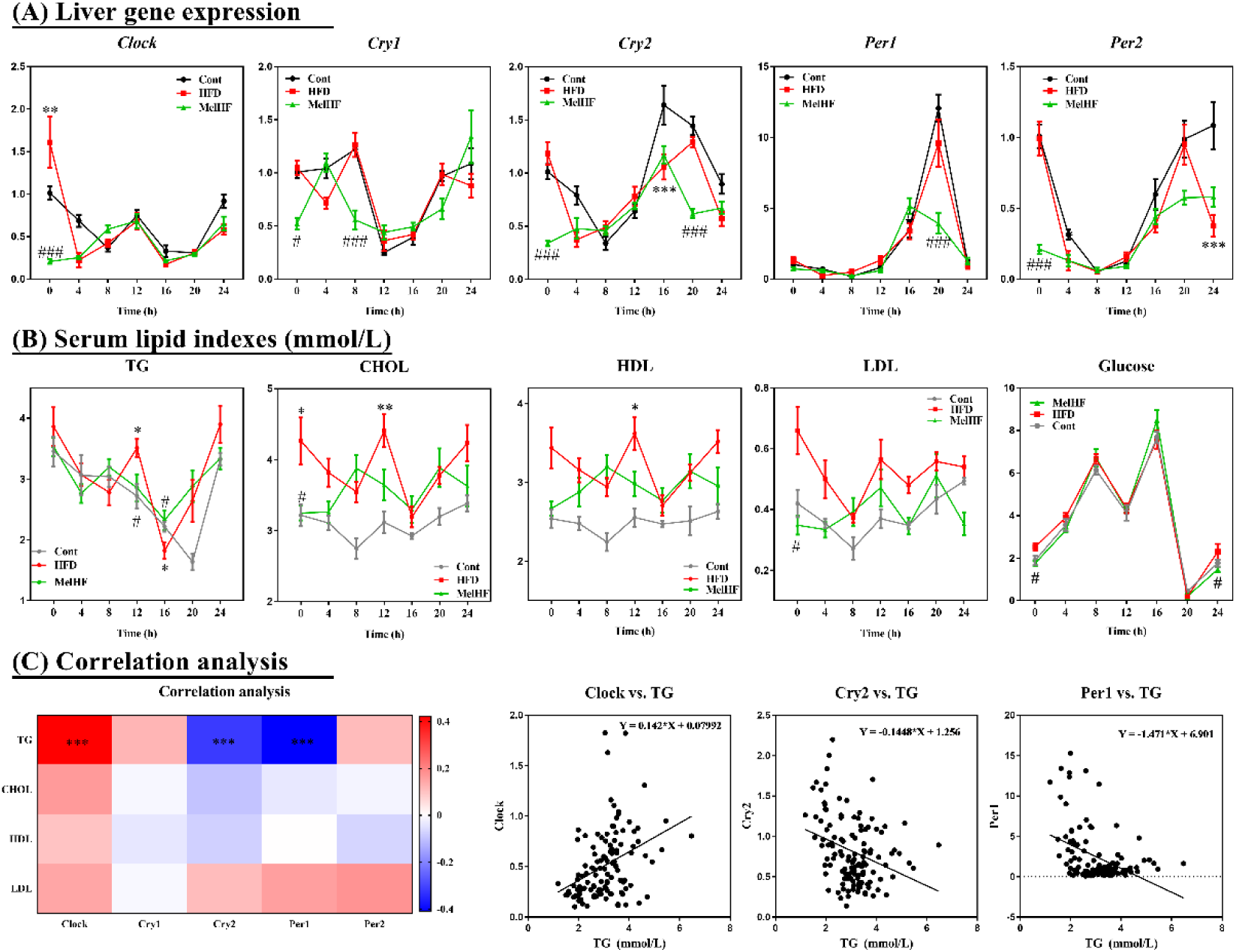
Effects of admistration of exogenous melatonin on diurnal rhythmicity of liver clock genes mRNA (*Clock*, *Cry1*, *Cry2*, *Per1*, and *Per2*) and serum lipid levels (TG, CHOL, HDL, LDL, and glucose) in HFD-fed mice. (A) Liver gene expression, (B) serum lipid levels, and (C) correlation analysis between circadian clock genes and serum lipid indexes. Gene expression was determined by real-time PCR analysis and relative gene pressions were normalized with β-actin. Values are presented as mean ± sem. Differences between groups were assessed by Bonferroni test and denoted as follows: ^*/#^P<0.05; ^**/##^P<0.01; ^***/###^P<0.001. * means difference between control and HFD groups at (A) and (B); # means difference between HFD and MelHF groups at (A) and (B); Correlation analysis was conducted by Pearson correlation analysis and the correlation coefficient was used for the heatmap: ***P<0.001.

**Table 1.**
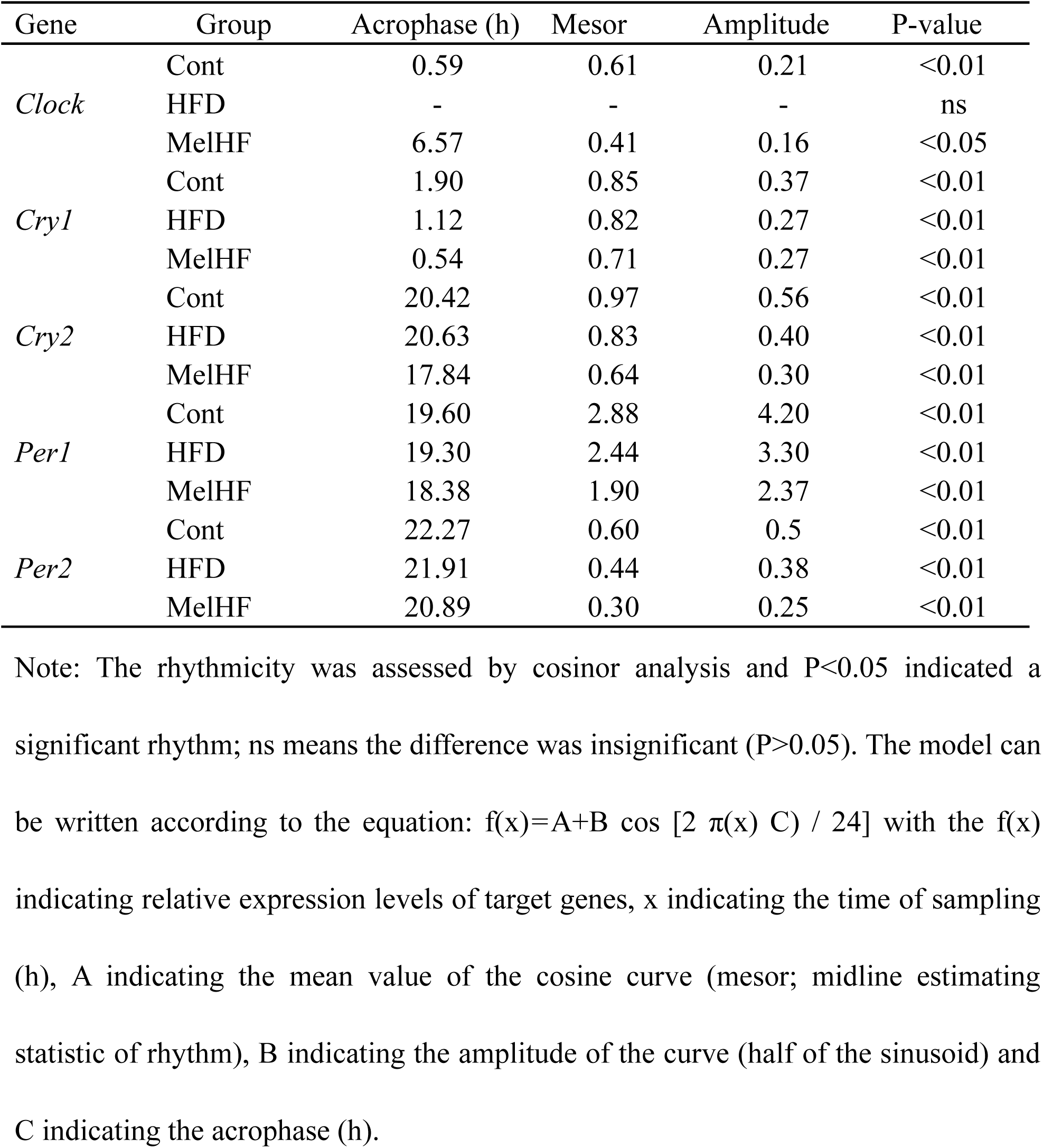
Mesor, amplitude and Acrophase of mRNA levels of clock genes in the liver of control, HFD, and MelHF mice.

| Gene | Group | Acrophase (h) | Mesor | Amplitude | P-value |
| --- | --- | --- | --- | --- | --- |
| *Clock* | Cont | 0.59 | 0.61 | 0.21 | <0.01 |
|  | HFD | - | - | - | ns |
|  | MelHF | 6.57 | 0.41 | 0.16 | <0.05 |
| *Cry1* | Cont | 1.90 | 0.85 | 0.37 | <0.01 |
|  | HFD | 1.12 | 0.82 | 0.27 | <0.01 |
|  | MelHF | 0.54 | 0.71 | 0.27 | <0.01 |
| *Cry2* | Cont | 20.42 | 0.97 | 0.56 | <0.01 |
|  | HFD | 20.63 | 0.83 | 0.40 | <0.01 |
|  | MelHF | 17.84 | 0.64 | 0.30 | <0.01 |
| *Per1* | Cont | 19.60 | 2.88 | 4.20 | <0.01 |
|  | HFD | 19.30 | 2.44 | 3.30 | <0.01 |
|  | MelHF | 18.38 | 1.90 | 2.37 | <0.01 |
| *Per2* | Cont | 22.27 | 0.60 | 0.5 | <0.01 |
|  | HFD | 21.91 | 0.44 | 0.38 | <0.01 |
|  | MelHF | 20.89 | 0.30 | 0.25 | <0.01 |

Note: The rhythmicity was assessed by cosinor analysis and P<0.05 indicated a significant rhythm; ns means the difference was insignificant (P>0.05). The model can be written according to the equation: $f(x)=A+B \cos [2 \pi(x) C] / 24$ with the f(x) indicating relative expression levels of target genes, x indicating the time of sampling (h), A indicating the mean value of the cosine curve (mesor; midline estimating statistic of rhythm), B indicating the amplitude of the curve (half of the sinusoid) and C indicating the acrophase (h).

### 3.3 Diurnal rhythms of serum lipid in response to HFD and exogenous melatonin

Next, we determined the diurnal patterns of serum lipid and glucose in the three experimental groups (Figure 2B; Table 2). Serum TG exhibited significant rhythmicity in control and MelHF mice (P<0.01), but not in HFD mice (P>0.05). Significant diurnal rhythm of LDL only occurred in control mice (P<0.01). Serum glucose exhibited rhythmicity in all the three groups (P<0.01). No daily rhythms were observed in the levels of serum CHOL and HDL (P>0.05). Despite rhythmicity, all lipid and glucose indexes were widely higher in HFD-fed mice, while the trends in the MelHF group were similar to the control subjects and the values were widely decreased compared with the HFD-fed mice at specific time points, as previously shown [17].

**Table 2.** Mesor, amplitude and Acrophase of serum lipid indexes.

| Item | Group | Acrophase (h) | Mesor | Amplitude | P-value |
| --- | --- | --- | --- | --- | --- |
| TG | Cont | 0.79 | 2.96 | 0.75 | <0.01 |
|  | HFD | - | - | - | ns |
|  | MelHF | 0.62 | 3.13 | 0.45 | <0.01 |
| CHOL | Cont | - | - | - | ns |
|  | HFD | - | - | - | ns |
|  | MelHF | - | - | - | ns |
| HDL | Cont | - | - | - | ns |
|  | HFD | - | - | - | ns |
|  | MelHF | - | - | - | ns |
| LDL | Cont | 23.2 | 0.39 | 0.06 | <0.01 |
|  | HFD | - | - | - | ns |
|  | MelHF | - | - | - | ns |
| Glucose | Cont | 11.56 | 3.56 | 2.87 | <0.01 |
|  | HFD | 10.47 | 3.72 | 2.93 | <0.01 |
|  | MelHF | 11.5 | 3.98 | 2.89 | <0.01 |

Note: The rhythmicity was assessed by cosinor analysis and $P < 0.05$ indicated a significant rhythm; ns means the difference was insignificant ( $P > 0.05$ ). The model can be written according to the equation: $f(x) = A + B \cos [2 \pi(x) C / 24]$ with the $f(x)$ indicating relative expression levels of target genes, $x$ indicating the time of sampling (h), $A$ indicating the mean value of the cosine curve (mesor; midline estimating statistic of rhythm), $B$ indicating the amplitude of the curve (half of the sinusoid) and $C$ indicating the acrophase (h).

To check if the serum lipid rhythmicity was associated with the liver expression of clock genes, we performed Pearson correlation analysis among serum lipid indexes and circadian clock genes (*Clock*, *Cry1*, *Cry2*, *Per1*, and *Per2*) (Figure 2C). Surprisingly, serum TG concentration was positively correlated with C*lock* expression but exhibited a negative correlation with the mRNA levels of *Cry2* and *Per1* (P<0.001). Together, the rhythmicity of lipid indexes widely existed in the blood and was markedly associated with clock genes expression, especially for TG concentration. The daily rhythm TG was impaired in the HFD-fed mice, which was markedly improved by administration of exogenous melatonin.

### 3.4 Effect of Melatonin on the diurnal rhythms of gut microbiota in HFD-fed mice

Gut microbiota has been identified as a key element involving in host circadian rhythms and itself also undergoes circadian oscillation, which is disturbed in HFD-fed mice or obesity model [23, 24, 42]. Our previous study demonstrated that melatonin treatment improved lipid metabolism by reprogramming gut microbiota in HFD-fed mice [17], thus we hypothesized that administration of exogenous melatonin will improve the daily rhythm of gut microbiota.

Mice were sacrificed every 4 h within 24 h period and metagenomic DNA was extracted from the cecal contents. Gut microbiota was tested by 16S rDNA sequencing and the compositions were similar to our previous study [17] that the largest phyla *Bacteroidetes* was reduced and *Firmicutes* abundance was increased in HFD-fed mice, while melatonin reversed these alterations (Figure 3A). *Firmicutes* exhibited a significant rhythmicity in control and MelHF mice (P<0.05), but not in HFD mice (P>0.05), while *Bacteroidetes* exhibited rhythmicity only in the control and HFD groups (P<0.05) (Figure 3C; Table 3). *Firmicutes* relative abundance peaked at 4:00 in the HFD group, but at 8:00 in the control and MelHF groups (Figure 3C). However, *Proteobacteria* and *Actinobacteria* failed to show a diurnal variation at the phylum (Figure 3C; Table 3).

**Figure 3.**
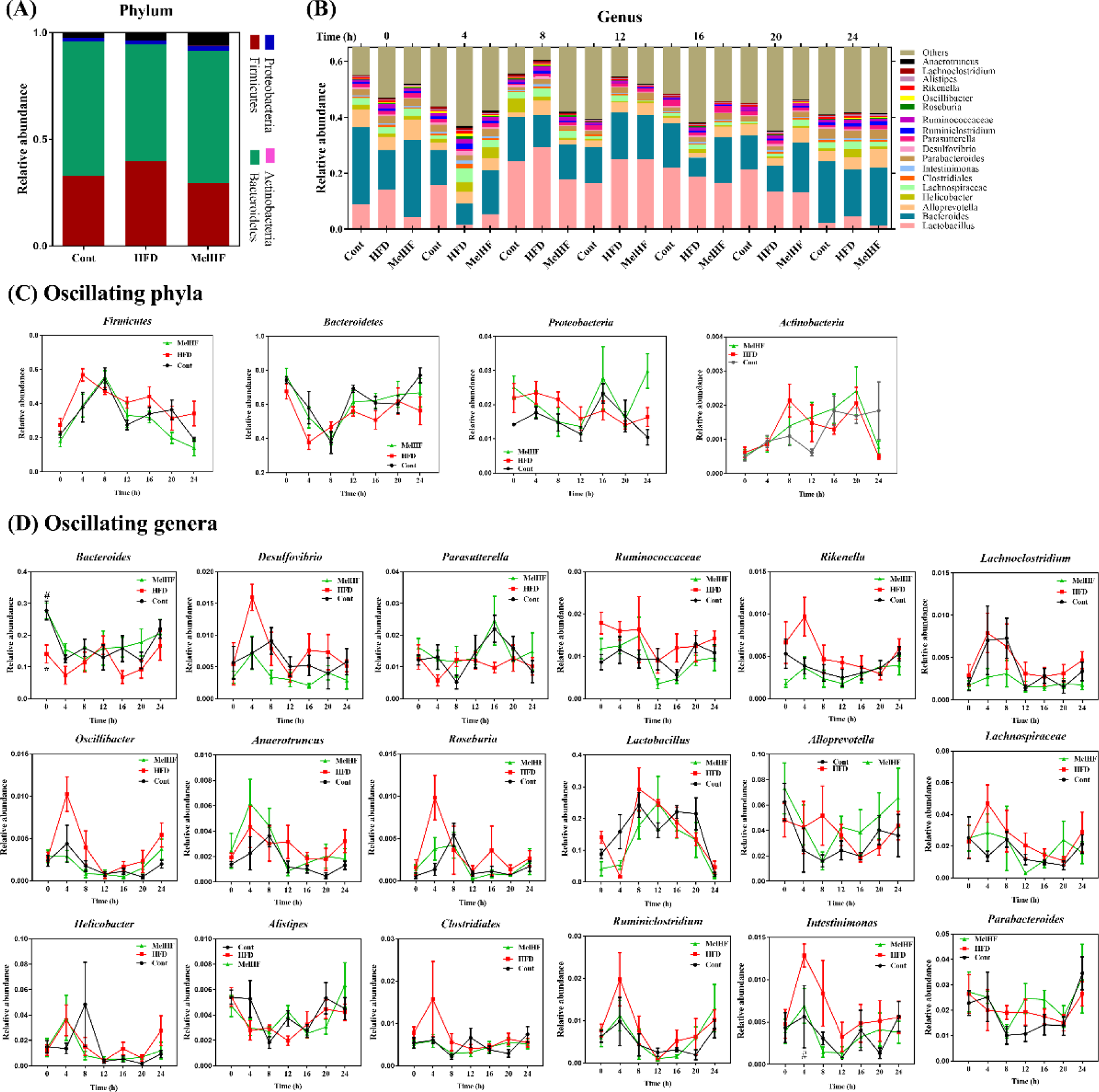
Admistration of exogenous melatonin improved the compositions and diurnal rhythmicity of gut microbiota in HFD-fed mice. (A) Microbiota compositions at the phylum level, (B) Microbiota compositions at the genus level, (C) Oscillating phyla, (D) oscillating genera. Values are presented as mean ± sem. Differences between groups were assessed by Bonferroni test and denoted as follows: ^*/#^P<0.05. * means difference between control and HFD groups; # means difference between HFD and MelHF groups.

**Table 3.**
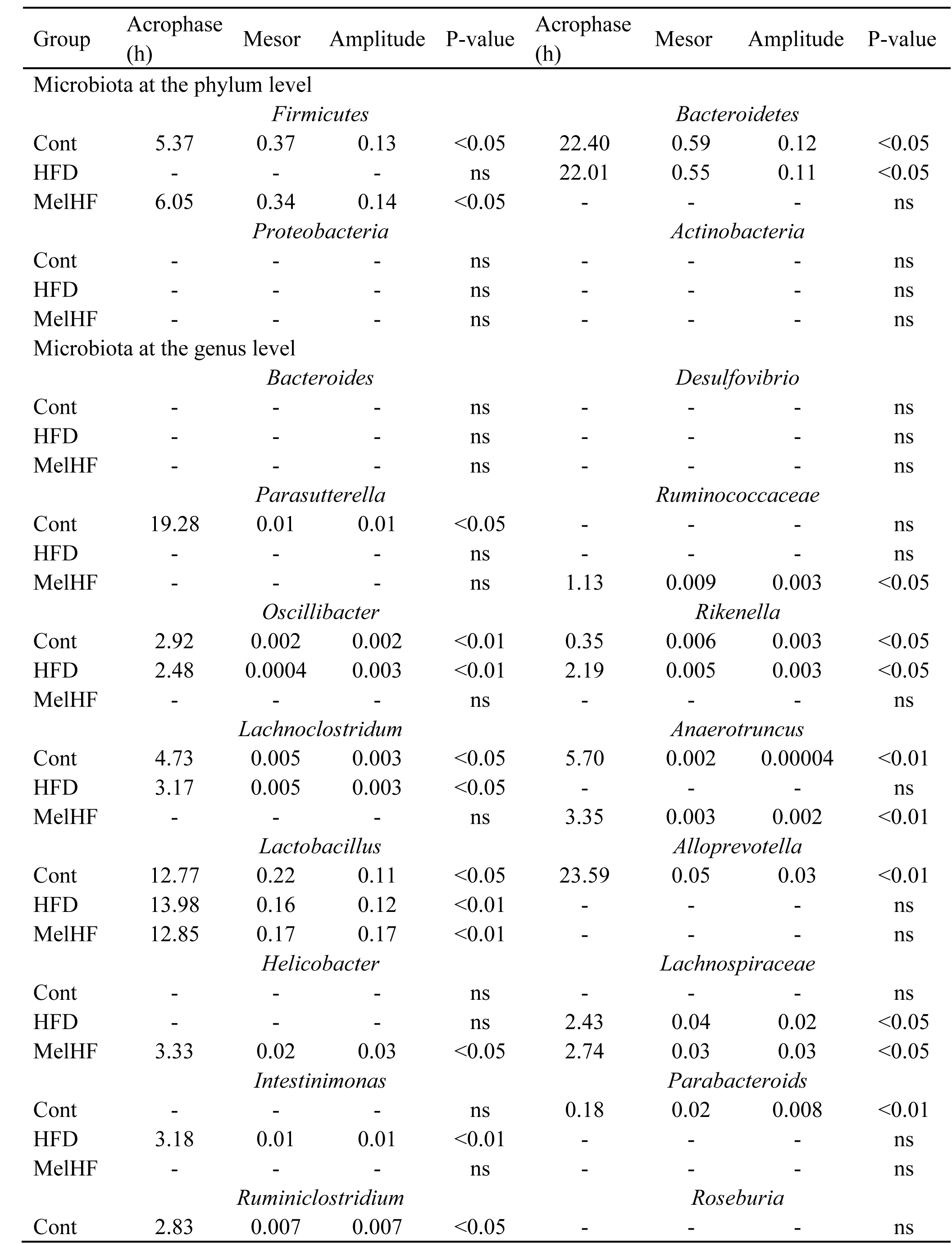
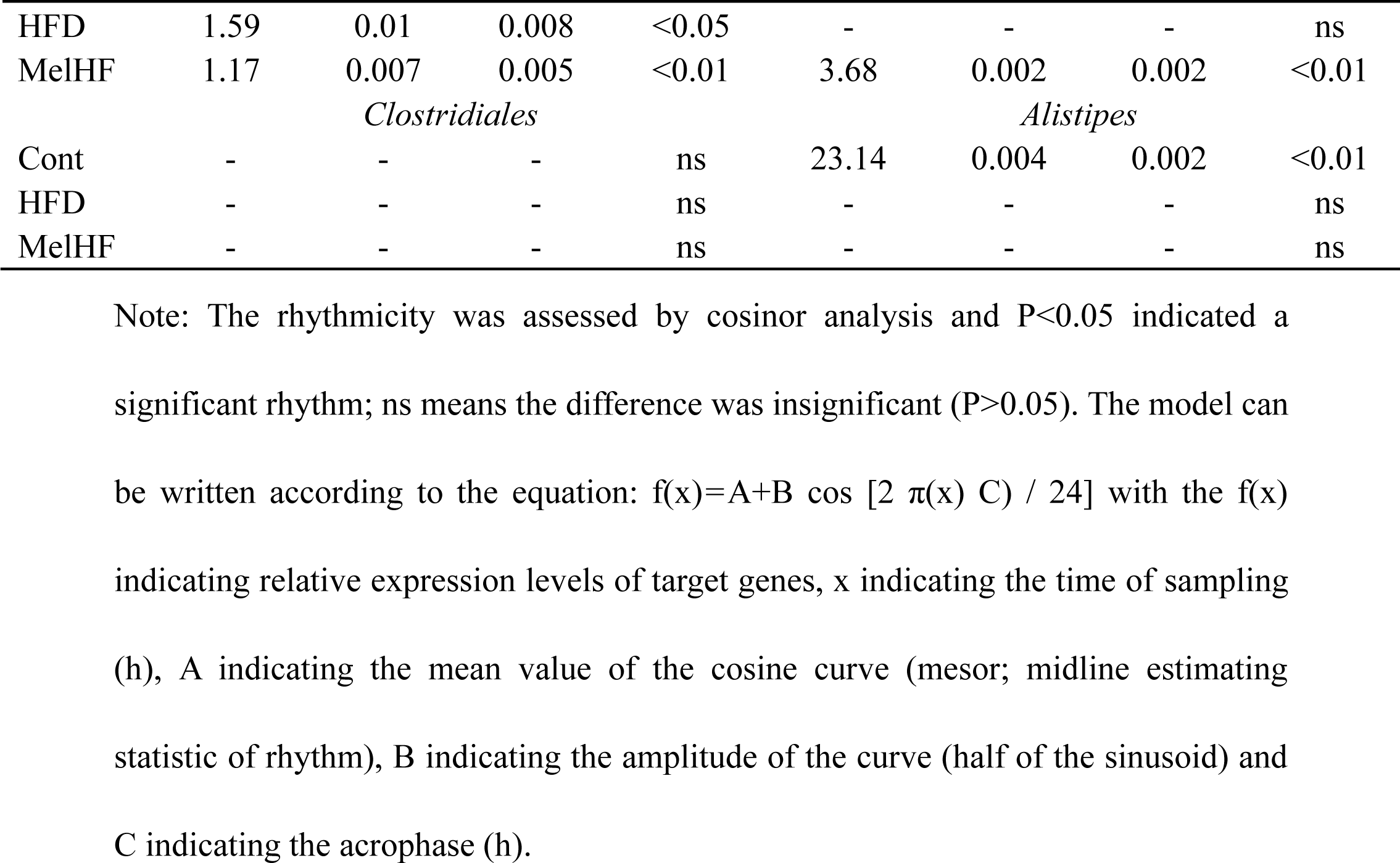
Mesor, amplitude and Acrophase of gut microbiota compositions.

At the genus level, 18 genera were mainly analyzed and most of them exhibited a marked daily rhythmicity expect for *Bacteroides*, *Desulfovibrio*, and *Clostridiales* (P>0.05) (Figure 3B and D; Table 3). *Parasutterella* (P<0.05), *Alloprevotella* (P<0.01), *Parabacteroids* (P<0.01), and *Alistipes* (P<0.01) were only rhythmic in the control group. *Intestinimonas* exhibited a significant rhythm only in HFD-fed mice (P<0.01). *Ruminococcaceae* (P<0.05), *Helicobacter* (P<0.05) and *Roseburia* (P<0.01) showed a daily rhythm only in the melatonin treated mice. *Oscillibacter*, *Rikenella*, and *Lachnoclostridum* were markedly cycled in the control and HFD groups (P<0.05), but not in the MelHF group (P>0.05). *Anaerotruncus* showed a diurnal pattern only in the control and MelHF groups (P<0.01), but not in HFD-fed mice (P>0.05). We also noticed that *Lachnospiraceae* was rhythmic in the HFD and MelHF groups (P<0.05), but not in the control group (P>0.05). In addition, *Lactobacillus* and *Ruminiclostridium* exhibited rhythmicity regardless of HFD and melatonin challenges (P<0.05).

Collectively, our data shown that most of the microbiota exhibited a daily variation and the rhythmicity of the microbiota was similar between control and MelHF groups, suggesting that the diurnal network of gut microbiota was affected by HFD and reversed, at least in part, by administration of exogenous melatonin (Additional file 2: Supplementary figure 1).

### 3.5 Genome prediction of microbial communities

Metabolism, genetic information, environmental information, cellular processes, human diseases, and organismal systems pathways were further annotated according to the microbiota compositions by Tax4Fun analysis (Figure 4A). Our data show that short-term HFD feeding markedly affected cell growth and death, endocrine and metabolic diseases, endocrine system, nervous system, immune system, and environmental adaptation (P<0.05), while administration of exogenous melatonin influenced lipid metabolism and terpenoids and polyketides (P<0.05). We then further analyzed the lipid metabolism (Figure 4B) and we identified eight pathways that mainly contributed to lipid metabolism-annotated genes, including lipid biosynthesis proteins, fatty acid biosynthesis, glycerophospholipid metabolism, glycerolipid metabolism, sphingolipid metabolism, fatty acid degradation, biosynthesis of unsaturated fatty acids, and synthesis and degradation of ketone bodies (Figure 4C).

**Figure 4.**
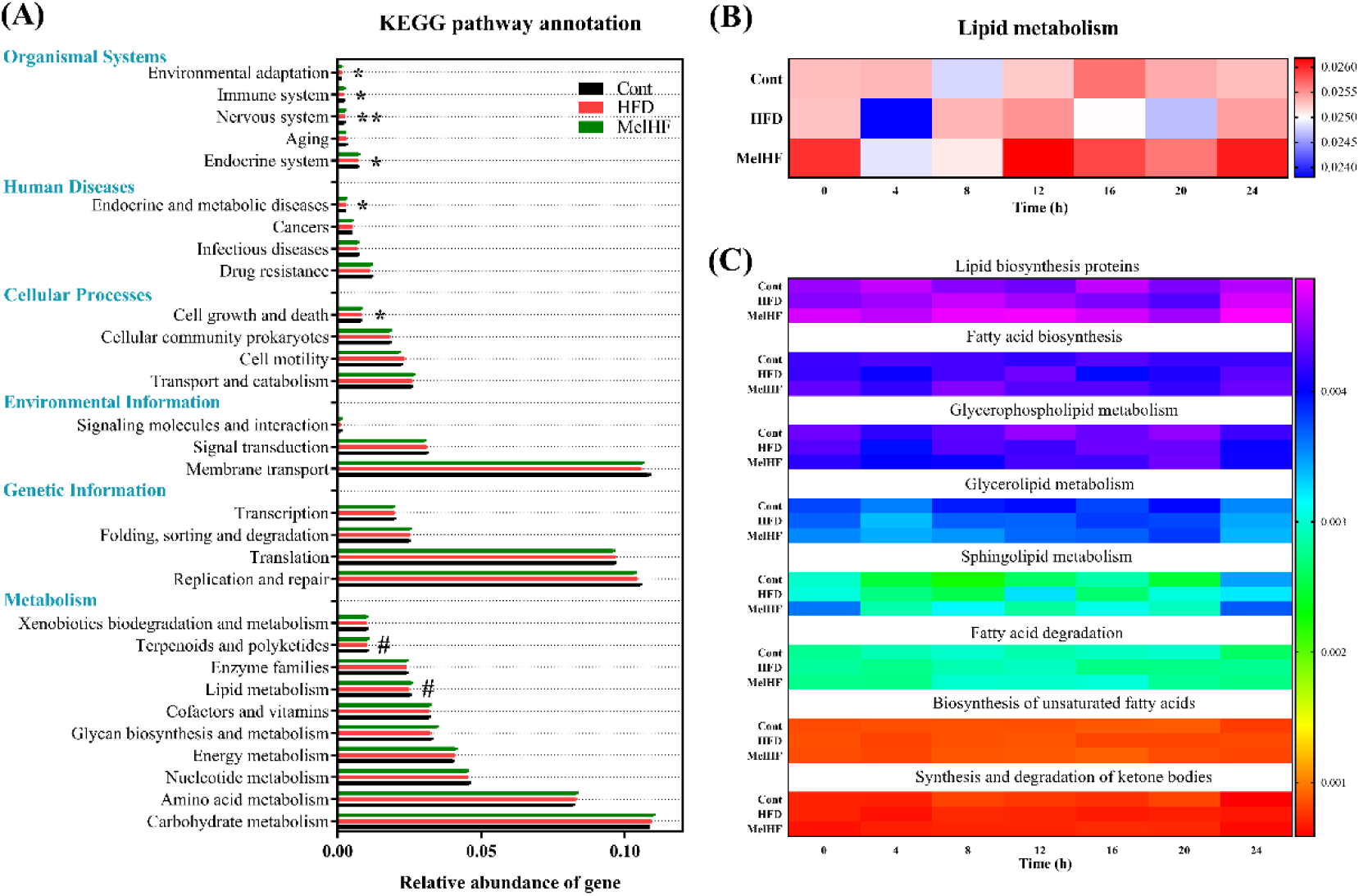
Predictive functional profiling of microbial communities by Tax4Fun analysis. (A) KEGG pathway annotations, (B) Lipid metabolism, and (C) detailed analysis of lipid metabolic pathways. Differences were assessed by Bonferroni test and denoted as follows: ^*/#^P<0.05; ^**/##^P<0.01. * means difference between control and HFD groups; # means difference between HFD and MelHF groups.

### 3.6 Gut microbiota correlated with clock genes and serum lipid levels

We then investigate whether gut microbiota also show an association with clock genes expression and serum lipid levels (Figure 5A). Four genera (*Alloprevotella*, *Parabacteroides*, *Rikenella*, and *Alistipes*) were identified to be positively correlated to *Clock* mRNA (P<0.05) (Figure 5B), while *Helicobacter*, *Lachnospiraceae*, and *Anaerotruncus* showed a negative association with *Per1* mRNA (P<0.05) (Figure 5C). Six genera exhibited high correlation with *Per2* mRNA, which positively correlated to *Alloprevotella* (P<0.05), *Rikenella* (P<0.01), and *Alistipes* (P<0.05) and negatively associated with *Lactobacillus* (P<0.05), *Helicobacter* (P<0.05), and *Anaerotruncus* (P<0.01) (Figure 5D). *Cry1* mRNA showed a negative correlation with *Lactobacillus* (P<0.05) but positively correlated with *Alloprevotella* (P<0.05), *Ruminococcaceae* (P<0.05), *Oscillibacter* (P<0.05), and *Rikenella* (P<0.01) (Figure 5E). Interestingly, *Helicobacter* (P<0.01), *Lachnospiraceae* (P<0.01), *Intestinimonas* (P<0.05), *Ruminiclostridium* (P<0.05), *Roseburia* (P<0.05), *Lachnoclostridium* (P<0.05), and *Anaerotruncus* (P<0.01) were found to be correlated to *Cry2* mRNA (Figure 5F). Together, fourteen genera were found to correlate with clock genes expression, these correlations were mostly positive with *Clock* and *Cry1* mRNA and negative *Cry2* and *Per1* mRNA levels.

**Figure 5.**
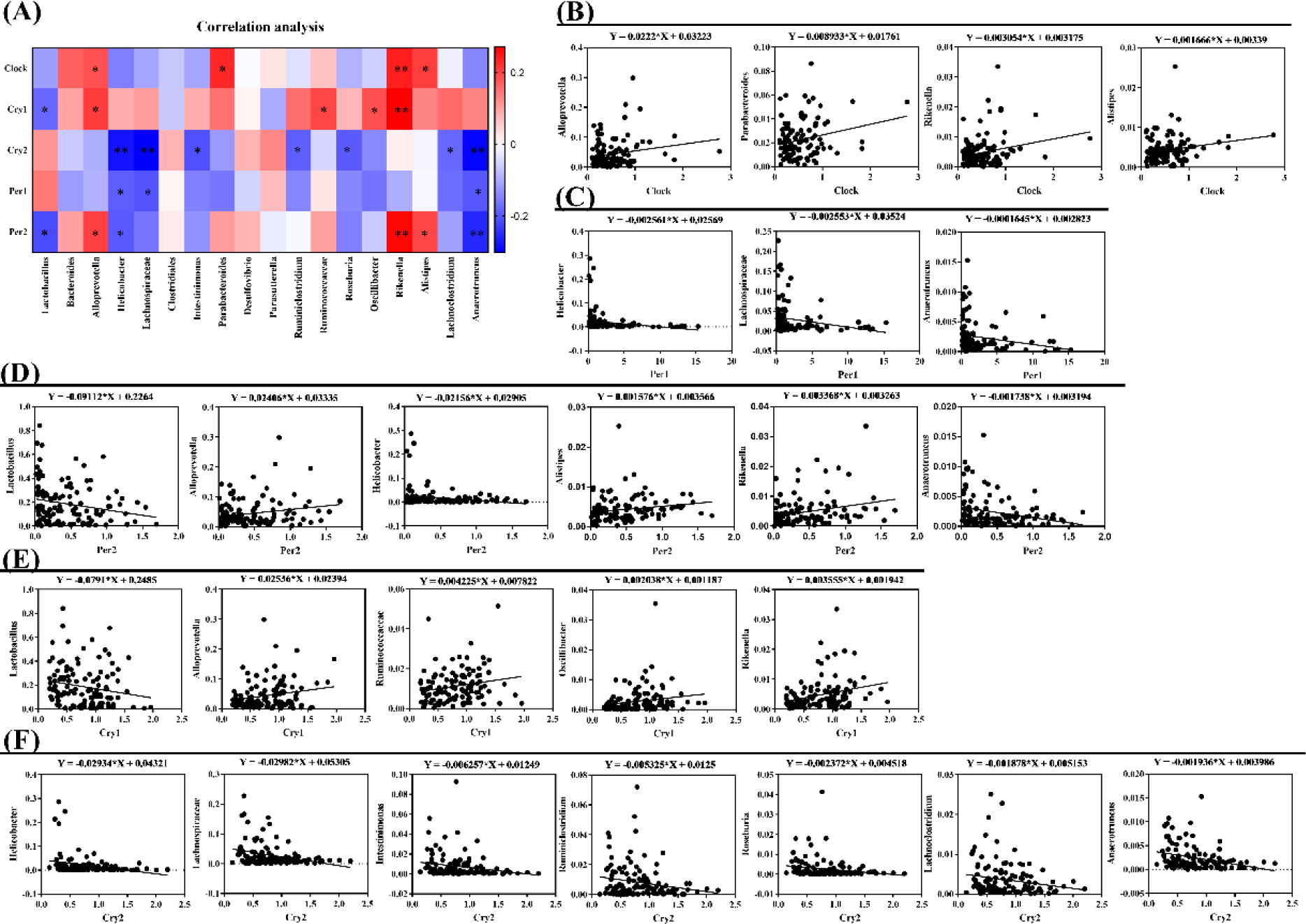
Correlation analysis between clock genes expression and gut microbiota at the genus level. (A) Overview of correlation analysis between circadian clock genes and gut microbiota at the genus level, (B) *Clock* gene correlated microbiota, (C) *Per1* gene correlated microbiota, (D) *Per2* gene correlated microbiota, (E) *Cry1* gene correlated microbiota, and (F) *Cry2* gene correlated microbiota. Correlation analysis was conducted by Pearson correlation analysis and the correlation coefficient was used for the heatmap: *P<0.05; **P<0.01.

Correlation analysis between serum lipid indexes and gut microbiota was further conducted and marked correlations were only noticed in TG and LDL concentrations (Figure 6A). Only one genus (*Lactobacillus*) was identified to be the negative correlated with TG (P<0.01), a positive correlation was found with *Bacteroides* (P<0.01), *Helicobacter* (P<0.01), *Parabacteroides* (P<0.01), *Ruminiclostridium* (P<0.05), *Rikenella* (P<0.05), and *Alistipes* (P<0.01) (Figure 6B). The relationship between LDL and *Rikenella* (P<0.05), *Alistipes* (P<0.05), and *Clostridiales* (P<0.01) exhibited a positive linear dependence (Figure 6C).

**Figure 6.**
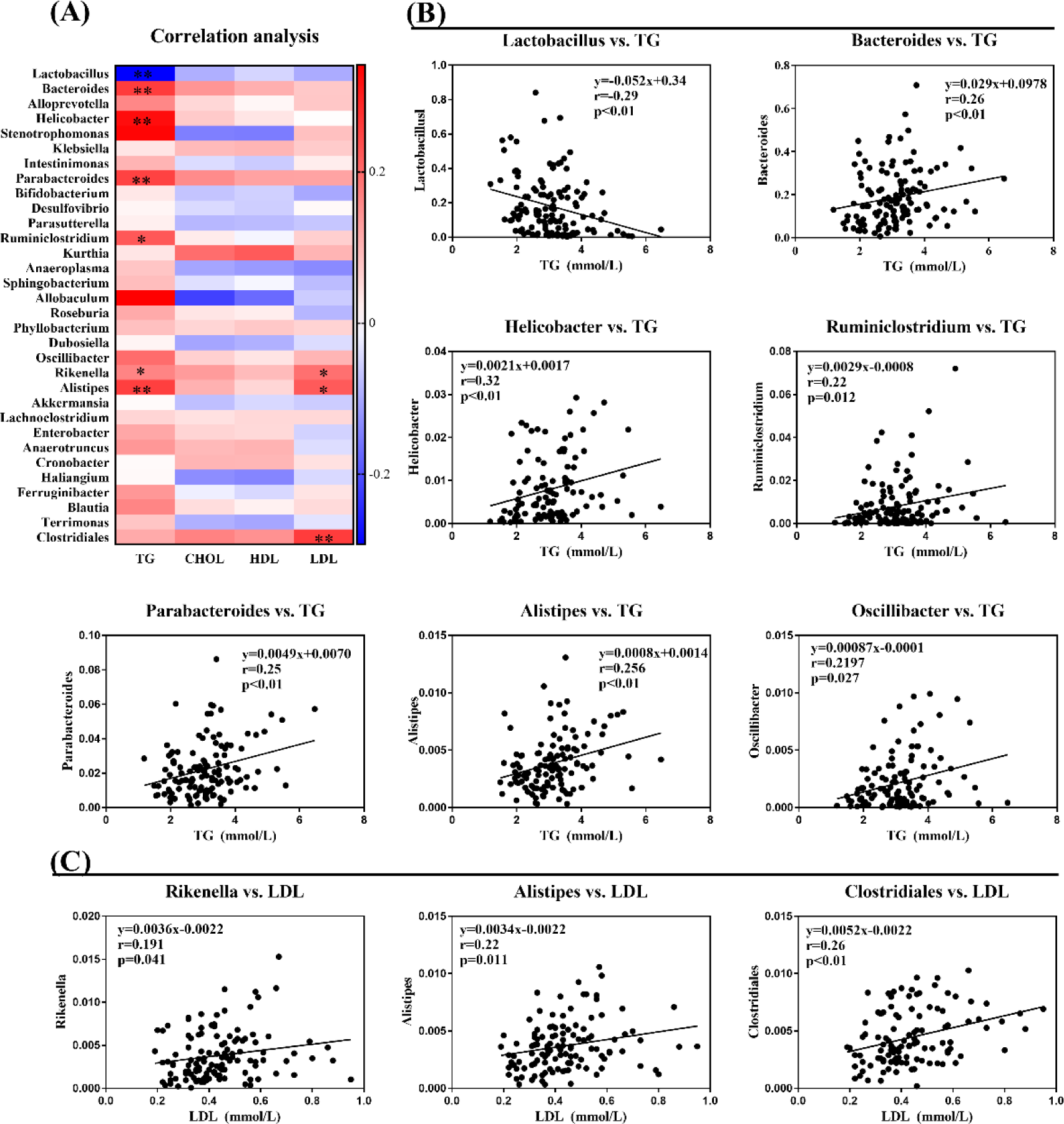
Correlation analysis between serum lipid levels and gut microbiota at the genus level. (A) Overview of correlation analysis between serum lipid indexes and gut microbiota at the genus level, (B) serum TG correlated microbiota, and (C) serum LDL correlated microbiota. Correlation analysis was conducted by Pearson correlation analysis: *P<0.05; **P<0.01. Correlation analysis was conducted by Pearson correlation analysis and the correlation coefficient was used for the heatmap: *P<0.05; **P<0.01.

### 3.7 Administration of exogenous melatonin at daytime or nighttime exhibited different effects on lipid accumulation and gut microbiota compositions

Administration of exogenous melatonin improved the daily rhythmicity in HFD-fed mice, so we further determined the effect of melatonin treatment at daytime or nighttime on lipid metabolism and gut microbiota. HFD-fed mice showed higher relative weight of subcutaneous inguinal fat, periuterine fat, perirenal fat, and total fat (P<0.001) (Figure 7A-E). Admistration of exogenous melatonin at daytime markedly reduced perirenal fat (P<0.05) and total fat (P<0.01) (Figure 7D and E), but nighttime-treatment of melatonin did not produce any significant effect (P>0.05) (Figure 7B-E). We also tested serum lipid indexes (Figure 7F-J) and the results showed that serum TG and bile acid concentrations were markedly reduced in the MelD group (P<0.05) but not in the MelN group (P>0.05).

**Figure 7.**
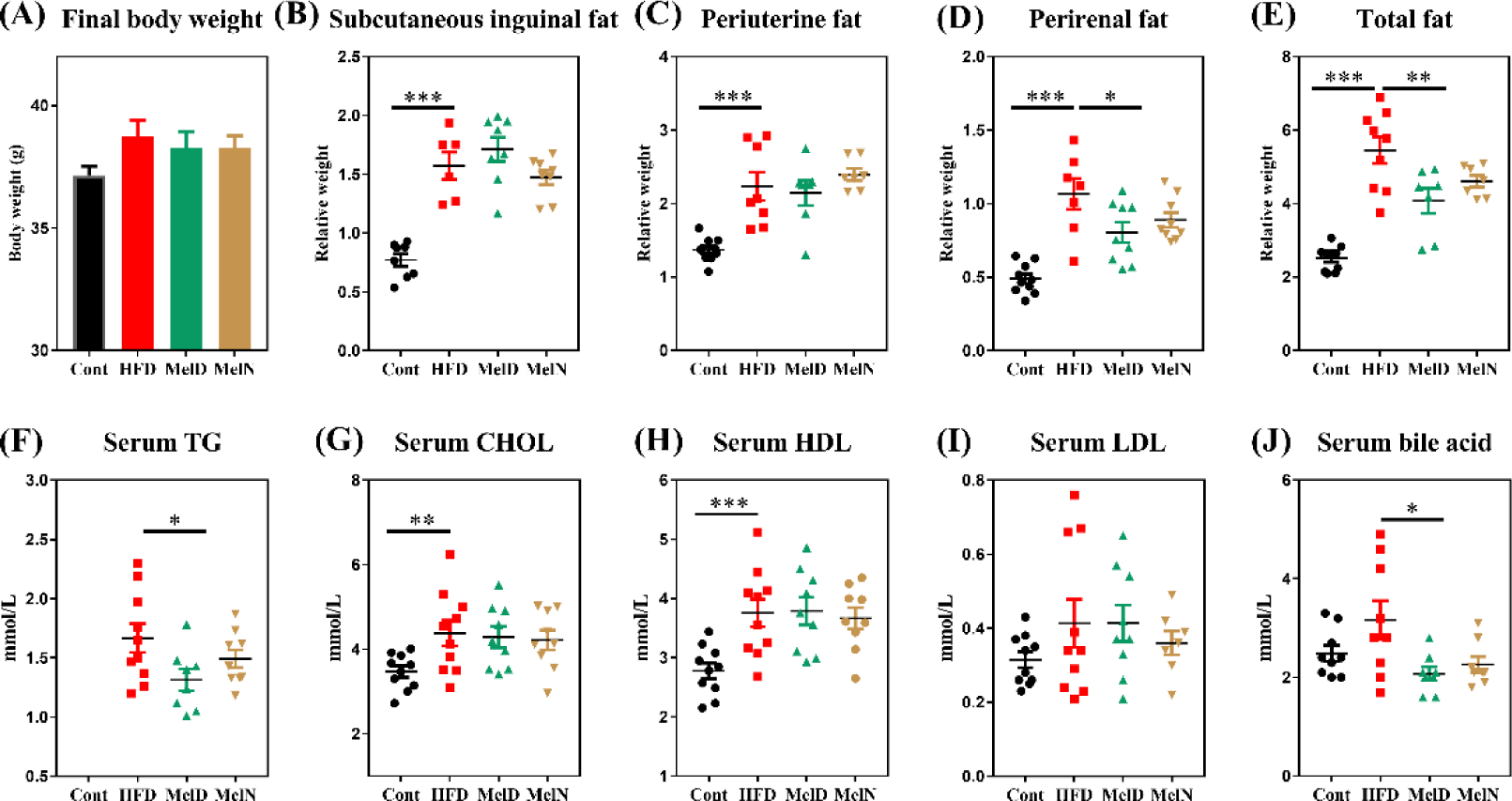
Melatonin treatment at daytime and nighttime exhibited different effects on lipid accumulation in HFD-fed mice. (A) Final body weight (g), (B) relative weight of subcutaneous inguinal fat, (C) relative weight of periuterine fat, (D) relative weight of perirenal fat, (E) relative weight of total fat, (F) serum TG concentration (mmol/L), (G) serum CHOL concentration (mmol/L), (F) serum HDL concentration (mmol/L), (F) serum LDL concentration (mmol/L), and (F) serum bile acid concentration (mmol/L). Values are presented as mean ± sem. Differences between groups were assessed by Bonferroni test and denoted as follows: *P<0.05; **P<0.01; ***P<0.001.

We then investigated gut microbiota compositions from HFD, MelD, and MelN groups using 16S rDNA. At the phylum level, both melatonin treatment at daytime or nighttime failed to alter gut microbiota compositions (Figure 8A). Interestingly, administration of exogenous melatonin during the nighttime significantly reduced the relative abundance of *Firmicutes* compared with the daytime treatment (P<0.05). At the genus level, *Lactobacillus*, *Intestinimonas*, and *Oscillibacter* were significantly affected by melatonin treatment during the day or the night (P<0.05) (Figure 8B).

**Figure 8.**
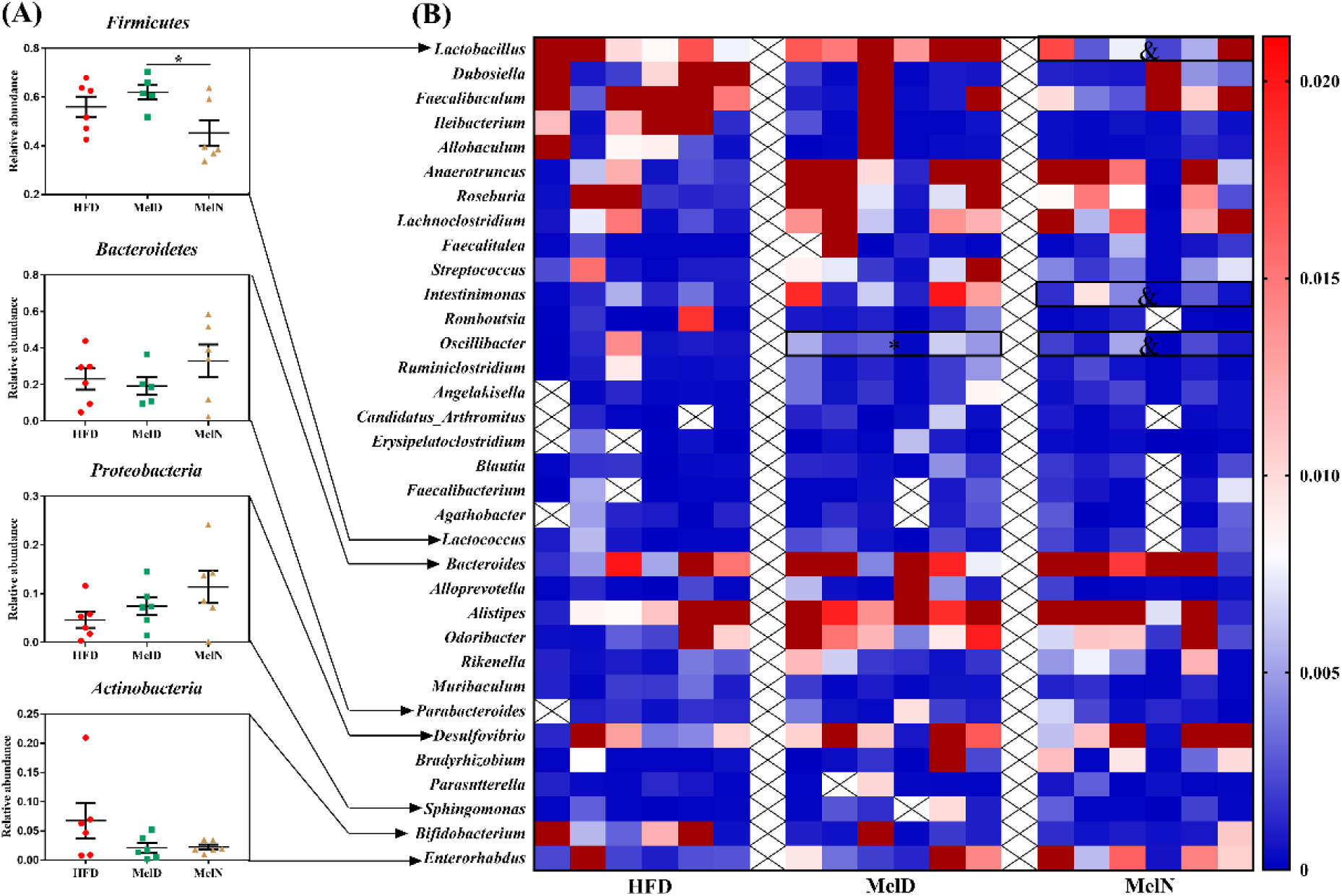
Melatonin treatment at daytime and nighttime exhibited different effects on gut microbiota compositions in HFD-fed mice. Microbiota at the phylum (A) and genus (B) levels. Values are presented as mean ± sem. Differences between groups were assessed by Bonferroni test and denoted as follows: ^*/&^P<0.05; * means difference was significant compared with HFD group at (B); & means difference was significant between MelD and MelN groups at (B).

### 3.8 Microbiota transplantation at different time of the day affected lipid metabolism in HFD-fed mice

We next transplanted fecal microbiota at two different time points (8:00 and 16:00) from control, HFD, and MelHF groups into antibiotics-treated mice to investigate the response to HFD feeding. Body weight was recorded and no significant difference was noticed between two time points (Figure 9A). Interestingly, the relative weight of subcutaneous inguinal fat at the MT-Cont group was affected by time at which the microbiota was transplanted (P<0.05). Overall, microbiota transplantation from HFD-fed mice tended to increase body weight, subcutaneous inguinal fat, perirenal fat, and periuterine fat, while microbiota transplantation from MT-MelHF group did not significantly affected the lipid accumulation (P>0.05) (Figure 9A-D).

**Figure 9.**
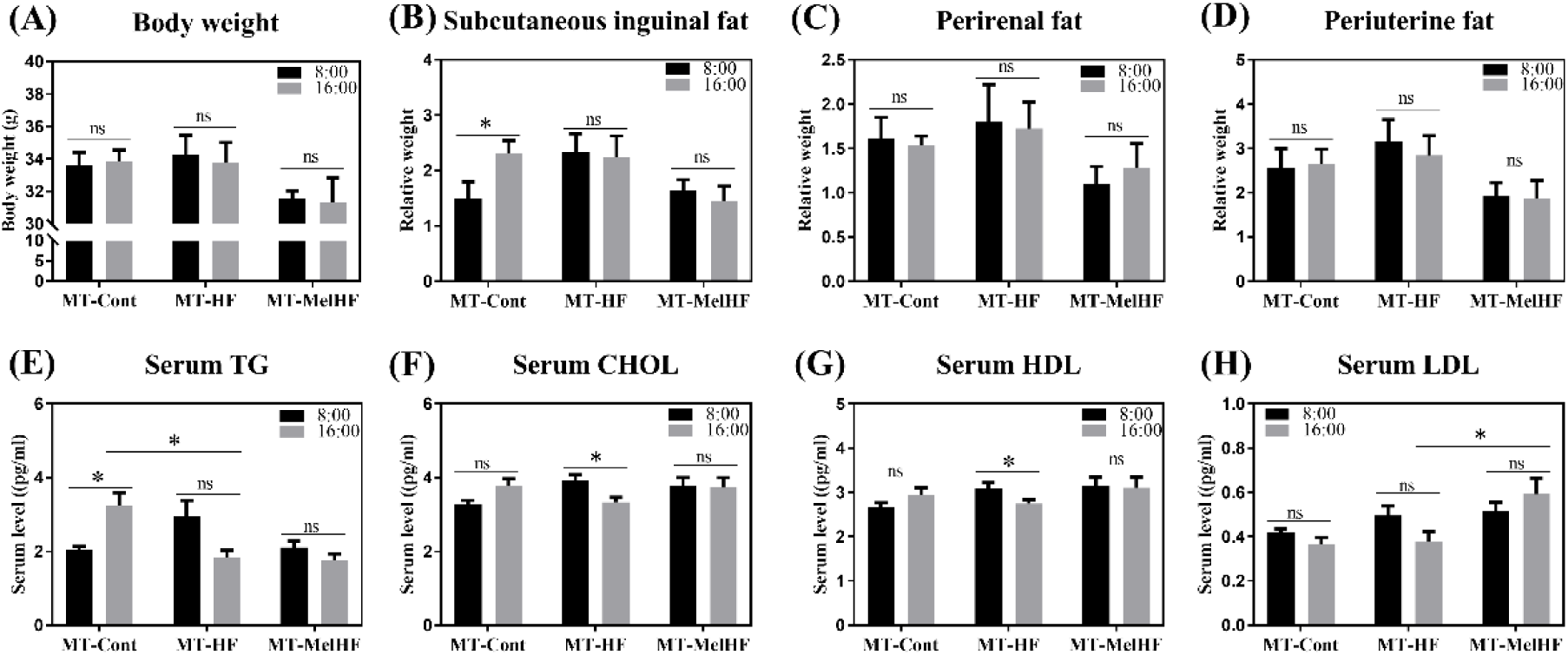
Microbiota transplantation at different time of the day affected lipid metabolism in HFD-fed mice. (A) Final body weight (g), (B) relative weight of subcutaneous inguinal fat, (C) relative weight of perirenal fat, (D) relative weight of periuterine fat, (E) serum TG concentration (mmol/L), (F) serum CHOL concentration (mmol/L), (G) serum HDL concentration (mmol/L), and (H) serum LDL concentration (mmol/L). The black bar means the fecal microbiota was transplanted at 8:00, while the gray bar at 16:00. Values are presented as mean ± sem. Differences between 8:00 and 16:00 in one group were assessed by student’s T test and multiple comparisons between groups (MT-Cont, MT-HF, and MT-MelHF) were analyzed by Bonferroni test and denoted as follows: *P<0.05; ^ns^P>0.05.

Similar to our previous study[17], microbiota transplantation at 8:00 from HFD group tended to enhance serum TG, CHOL, and HDL concentrations, which were slightly reversed in MT-MelHF group (Figure 9E-G). Conversely, serum TG concentration was reduced in MT-HF group (P<0.05) (Figure 9E) and LDL was higher in MT-MelHF group (P<0.05) (Figure 9H) at 16:00 microbiota transplanted mice. Notably, microbiota transplantation from control subjects at 8:00 tended to enhance serum CHOL and HDL (P>0.05) (Figure 9F and G) and significantly increased TG concentrations (P<0.05) (Figure 9E) compared with the 16:00 microbiota transplantation (Figure 9B). However, serum CHOL and and HDL levels were lower at 16:00 than these at 8:00 of HFD-derived microbiota transplantation (P<0.05) (Figure 9E-G). No difference was noticed between two time points in the MT-MelHF group.

## 4. Discussion

We have previously shown that admistration of exogenous melatonin improved HFD-induced lipid metabolic disorder by reversing gut microbiota compositions, especially for the relative abundances of *Firmicutes* and *Bacteroidetes* [17]. Here we have further confirmed that melatonin may reverse gut microbiota compositions in HFD-fed mice and gut microbiota are highly correlated to circadian clock genes and serum lipid indexes.

Ddiurnal rhythms and metabolism are tightly linked and obesity leads to profound reorganization of circadian system, leading remodeling of the coordinated oscillations between coherent transcripts and metabolites [43]. For example, 38 metabolites and 654 transcripts are identified to be oscillating only in HFD-fed animals and a majority of oscillations are clock-dependent [43, 44]. In this study, circadian clock genes (*Clock*, *Cry1*, *Cry2*, *Per1*, and *Per2*) and serum TG, LDL, and glucose concentrations exhibit a daily rhythmicity, which is similar to previous studies that most of circadian genes were rhythmic in the livers [45]. Interestingly, *Clock* and TG only cycled in the control and MelHF groups, but not in the HFD-fed mice, indicating that daily rhythmicity was impaired by short-term HFD feeding and admistration of exogenous melatonin partially rescued the daily rhythmicity in HFD-feed mice. Strikingly, serum TG concentration was positively correlated with C*lock* mRNA and negatively correlated with *Cry2* and *Per1* mRNA levels.

Compelling experimental evidence has shown a marked difference of gut microbiota between obese and lean subjects [46–48], here we further investigated the correlation between the microbiota (at the genus level) and the circadian clock genes and serum lipid levels. Fourteen genera showed a significant correlation between the clock genes expression. Positive correlations were observed with *Clock* and *Cry1* mRNA levels and negative correlations were observed with *Cry2* and *Per1*. Notably, *Alloprevotella* and *Rikenella* were found to be associated with *Clock*, *Cry1*, and *Per2*, whereas *Helicobacter* and *Anaerotruncus* correlated with the *Cry2*, *Per1*, and *Per2*. Previous studies have reported that germ-free mice show a reduced amplitude of clock gene expression in both central and peripheral tissues even in the presence of light-dark signals [23]. Together, our data may further indicate that the diurnal variations of clock genes may be governed, at least in part, by the gut microbiota. In addition, *Lactobacillus*, *Bacteroides*, *Helicobacter*, *Parabacteroides*, *Ruminiclostridium*, *Rikenella*, and *Alistipes* correlate to serum TG, and *Rikenella*, *Alistipes*, and *Clostridiales* are highly associated with LDL concentration. Among them, *Lactobacillus* has been extensively studied to involve in lipid accumulation [49–51], which is markedly enhanced in HFD-fed mice and reversed by administration of melatonin[17]. Our previous study indicates that *Bacteroides* and *Alistipes*-derived acetic acid targete host lipid metabolism [17], which is further corroborated by the current data that both *Bacteroides* and *Alistipes* are markedly associated with serum TG or LDL.

Microbiota analysis within 24 h period further confirms that admistartion of melatonin reverses gut microbiota compositions, especially for the relative abundances of *Firmicutes* and *Bacteroidetes* [16, 17]. In addition, we have also shown that the most gut microbiota exhibit daily cyclical variation in a variety of dietary and melatonin treatments [22–24, 42]. However, the diurnal variations of the gut microbiota are highly variable. For example, *Firmicutes* only cycled in control and MelHF mice (P<0.05), but not in HFD mice (P>0.05). Also, *Firmicutes* of HFD-fed mice peaks at 4:00 and markedly differentiates with control and MelHF groups, in which *Firmicutes* reachs the high point at 8:00. Conversely, HFD mice advance the low point of *Bacteroidetes* at 4: compared with 8:00 of the valley point in the control and MelHF groups. At the genus level, we also show that most genera oscillates within 24 h period and the cosine curves of microbiota are similar between control and MelHF groups, suggesting that the daily rhythm of gut microbiota is driven by HFD and reversed by melatonin administration. Microbiota rhythms have been indicated to represent a potential mechanism by which the gut microbiota affected host metabolism [22]. Using germ-free animal model, Thaiss *et al.* found that microbiota deficiency leads to a temporal reorganization of metabolic pathways evidenced by reduction of chromatin and transcript oscillations and a massive gain of de novo oscillations [52]. Together, our new data support the hypothesis that admistration of exogenous melatonin improves the diurnal rhythmicity of gut microbiota, which further regulates host metabolism.

Another important finding from the present study is that melatonin treatment at daytime and nighttime exhibit different responses in gut microbiota and HFD-induced lipid dysmetabolism, thus indirectly unveiling the role of gut microbiota rhythmicity in melatonin-mediated lipid metabolism. Our data show that melatonin treatment at daytime exhibits higher resistance to HFD-induced lipid dysmetabolism, indicating that duration of melatonin exposure is more important in lipid response. The reason may be associated with the secretory mechanism that melatonin mainly secrets at nighttime and melatonin treatment at daytime also maintains a higher abundance of melatonin, providing a sustained exposure of melatonin to host metabolism [53]. Intriguingly, we observed that *Lactobacillus*, *Intestinimonas*, and *Oscillibacter* responded to HFD and to the administration of melatonin at the different times of the day. Our previous study shows that *Lactobacillus* is increased in HFD-fed mice, which is reversed by administration of melatonin [17]. Similarly, the relative abundance of *Oscillibacter* is widely increased in HFD-fed mice [54, 55], indicating a potential role of *Lactobacillus*, *Intestinimonas*, and *Oscillibacter* in melatonin-mediated lipid metabolic response. Microbiota transplantation from different groups at different times show different susceptibility to HFD-induced lipid dysmetabolism, which further demonstrates the diurnal rhythmicity of gut microbiota. Notably, serum lipid indexes show a marked difference between two time points of microbiota transplantation from control and HFD mice but not from melatonin treated animals, indicating that the diurnal alteration of gut microbiota is affected by melatonin treatment.

## 5. Conclusion

In conclusion, our results show most gut microbiota exhibits daily rhythm and is highly correlated to clock genes expression and serum lipid levels. Melatonin improves the diurnal patterns of gut microbiota in HFD-fed mice, which is further confirmed by the administration of exogenous melatonin during the daytime or nighttime and microbiota transplantation. Melatonin treatment at daytime exhibited higher resistance to HFD-induced lipid dysmetabolism. Microbiota transplantation at the early in the morning or in the late afternoon also shows diverse responses to HFD. Thus, we conclude that most gut microbiota cycles within 24 h period and the rhythm is disturbed by HFD exposure. Melatonin treatment further improves diurnal patterns of gut microbial structure and function, which may further mediate host lipid metabolism in HFD-fed mice.

## Acknowledgements

We are grateful to the Public Service Technology Center, Institute of Subtropical Agriculture, Chinese Academy of Sciences for technical support. Many thanks also to the editors and reviewers for the painstaking care taken in helping improve the clarity of the manuscript.

## Availability of data and materials

Raw sequences are available in the NCBI Sequence Read Archive with the accession numbers (SAMN11246274-PRJNA528844 and SAMN11245315-PRJNA528812).

## Authors’ contribution

JY, TL, and YY designed the study. JY, YL, and HH conducted this study. JY, YL, and HH participated in samples collection and data analysis. KB conducted the cosinor analysis. JY drafted the manuscript. GL, PB, XW, XG, RJ, GQ, WK, BE, and XH given suggestions for this study. JY and GT revised the manuscript. All authors read and approved the final manuscript.

## Additional files

**Additional file 1: Supplementary table 1** Primers used in this study. (word 12.7kb)

**Additional file 2: Supplementary figure 1** Microbiota networks were markedly altered in the control, HFD, and MelHF groups. (TIF 4360kb)

